# Four Decades of Genomic Stability and Adaptive Divergence in *Xanthomonas arboricola* pv. *pruni*-Infecting Phages: Defining *Duraznoxanthovirus arenicola* and Its Evolutionary Framework

**DOI:** 10.1101/2025.11.25.690537

**Authors:** Katherine M. D’Amico-Willman, Prasanna Joglekar, Dann Turner, Meaghan Flaherty, David F. Ritchie, Alejandra I. Huerta

## Abstract

Bacteriophages (phages) are abundant and ecologically significant, yet their diversity and roles in plant-associated ecosystems remain poorly understood, limiting their application in sustainable disease management. To address this gap, we characterized 15 phages infecting *Xanthomonas arboricola* pv. *pruni*, the causal agent of bacterial spot on peach, has been isolated for over four decades from North Carolina orchards. Comparative genomic and phylogenetic analyses revealed two temporally distinct clades with >95% nucleotide identity and 63 conserved core genes, forming a new genus and species, *Duraznoxanthovirus arenicola*. These findings challenge assumptions of pervasive genomic mosaicism, highlighting remarkable genomic stability alongside localized variability in accessory loci. Beyond genus-level characterization, our analyses support a broader taxonomic restructuring within the family *Anamaviridae*, introducing a new subfamily (*Terravirinae*) and two new genera (*Duraznoxanthovirus* and *Ralstopathovirus*). This work provides the first family-level framework for phages exclusively infecting plant-associated bacteria, offering evolutionary insights and a foundation for ecological studies and management strategies.

## Introduction

Bacteriophages (phages) are ubiquitous viruses that infect bacteria, playing key ecological roles in microbial assemblages and influencing bacterial evolution. It is estimated that for every bacterial cell on earth, there are 10 phage particles [1]. Despite their abundance and ecological significance, the composition and functional roles of phage populations in plant-associated ecosystems remain largely unexplored [2]. This knowledge gap limits the potential of phage biology to manage bacterial plant diseases, particularly in agricultural settings [3]. To harness the biotechnological potential of phages through their lytic and lysogenic life cycles, it is essential to characterize their diversity and abundance and to understand their ecological impact in both managed and natural plant-associated ecosystems.

As of April 14, 2025, the InPhared database, which contains 34,076 publicly available, curated phage genomes from GenBank, includes only a limited number of genomes for phages that infect bacterial species within agriculturally important genera: 484 phages virulent on *Ralstonia*, 144 virulent on *Pectobacterium*, 53 virulent on *Dickeya*, 23 virulent on *Agrobacterium*, 264 virulent on *Erwinia*, and 200 virulent on *Xanthomonas* [4]. Within the genus *Xanthomonas*, there are 35 genomes for phages virulent on *X. campestris*, 36 virulent on *X. arboricola*, 34 virulent on *X. oryzae*, 15 virulent on *X. citri*, 10 virulent on *X. euvesicatoria*, 11 virulent on *X. translucens*, six virulent on *X. albilineans*, six virulent on *X. vesicatoria*, five virulent on *X. perforans*, two virulent on *X. fragariae*, and one each virulent on *X. hortorum* and *X. vesicola*, with an additional 38 genomes virulent on unclassified *Xanthomonas* strains. Notably, within the *X. arboricola* species, there are six genomes for phages virulent on *X. arboricola* pv. *pruni*, 23 for pv. *juglandis*, and seven phages infect both pvs. *corylina* and *juglandis*. This uneven representation across bacterial genera, species, and pathovars within agriculturally relevant bacterial taxa highlights knowledge gaps and opportunities for research in phage ecology and biotechnological applications.

*Xanthomonas arboricola* pv. *pruni* (Xap) causes bacterial spot on *Prunus* spp., including peach, nectarine, and almond [5]. The pathogen overwinters in plant tissues, becoming active in spring under favorable conditions, and spreads via rain, wind, and agronomic practices, infecting through natural openings to colonize the mesophyll to cause lesions, spots, and necrosis [6]. While phage applications have shown success in clinical and industrial settings, their effectiveness in field conditions remains inconsistent, including for Xap, where variability is likely driven by biotic and abiotic factors [3, 7, 8]. Peach’s perennial growth, clonal propagation, long production cycles, and wide geographic distribution make it an excellent model for studying phage diversity and phage-bacteria coevolution over time. However, knowledge of Xap-infecting phages in peach orchards is limited aside from the recent study by D’Amico-Willman et al (2025), which reported six phages (Xapϕ1-Xapϕ6) isolated over four decades from the same North Carolina orchard [9]. These phages cluster within the *Kantovirinae* subfamily but do not fit into any existing genus or species, highlighting the need for deeper taxonomic and ecological exploration of these Xap-virulent phages [9].

Phage taxonomy, once grounded in morphology and structural traits observed via electron microscopy, has undergone a paradigm shift driven by advances in rapid, cost-effective sequencing technologies [10, 11]. Modern classification now relies on genomic data, enabling deeper evolutionary insights into phage diversity and population structure, but also introducing challenges: many phages remain unclassified under the new framework [12, 13]. Furthermore, the majority of phages classified are virulent against human bacterial pathogens [12]. This bias limits our understanding of phage diversity in plant-associated ecosystems, highlighting the need for more comprehensive genomic characterization across plant-associated microbiomes.

Our program aims to understand how geographic distribution and temporal dynamics shape phage and bacterial host diversity across managed and natural plant-associated ecosystems. Accurate classification and naming are central to this effort, particularly in plant-associated systems where phage diversity remains poorly characterized. To advance this goal, we isolated, purified, and sequenced nine phages virulent on Xap. Comparative genomic analysis with 36 related phages in the recently described family *Anamaviridae* revealed two new genera and species: “*Duraznoxanthovirus arenicola*” [*durazno* = peach; *xantho* = *Xanthomonas*; *arenicola* = sand-dweller, referencing North Carolina’s Sandhills peach region] and “*Ralstopathovirus humicola*” [*Ralsto* = *Ralstonia*; *patho* = pathogen; *humicola* = soil-dweller] [14]. We also propose a new subfamily within the recently delineated phage family *Anamaviridae*, that we name “*Terravirinae*” [*terra* = soil], which includes the genera *Ralstopathovirus* and *Naesvirus* [14]. A formal taxonomic proposal for these taxa will be submitted to the ICTV; these names are used throughout the manuscript.

## Materials and Methods

### Bacterial and phage culture conditions

The bacterial host used as phage bait in this study was Xap strain Xcp1 [15]. The strain was routinely cultured on solid sucrose peptone agar (SPA) and in sucrose peptone broth at 28°C unless otherwise stated [16, 17]. Water agar (0.5% agar in sterile deionized water [SDW]) was used for double-layer overlay plate assays [18]. A 100 μL aliquot of log-growth phase stock inoculum was used to seed the top layer of double-layer overlays as described in D’Amico-Willman et al (2025). Phage isolates were routinely stored in Phage Buffer (PB) without glycerol (10 mM Tris [pH 7.5]; 10 mM MgSO^4^; 68 mM NaCl; 1 mM CaCl^2^) at 4°C (https://discoveryguide.seaphages.org<u>^/^</u>).

### Phage isolation

Peach leaves symptomatic of bacterial spot were collected from different cultivars, tissue types, and dates, at the Sandhills Research Station, Jackson Springs, NC, USA (Table 1). To isolate phages, symptomatic tissue was cut into strips with a sterile blade and incubated with approximately 10 mL of SDW in a 50-mL conical tube at room temperature overnight. After incubation, a 1-mL aliquot was treated with 8% chloroform (v/v) to inactivate any living bacteria. Double-layer overlay plates were prepared as described by D’Amico-Willman et al. (2025). A 10-µL aliquot of the chloroform-treated supernatant was spotted onto the Xcp1-seeded top agar. After 24 h at 28 °C, one plaque per sample was selected for subsequent phage purification and characterization. This was achieved by tracing the circumference of a plaque with a sterile toothpick, resuspending it in 1 mL of PB, and treating it with 8% chloroform (v/v). To further purify, a 20-µL aliquot was spotted onto a freshly prepared Xcp1-seeded top agar and streaked to form single plaques. This was repeated three times to ensure phage purification. Finally, 20 µL of the phage solution was spread onto an Xcp1-seeded top agar plate and incubated at 28°C for 24 h. The plate was then flooded with 3 mL of PB and incubated at room temperature for an additional 24 h. The resulting supernatant was aspirated from the plate surface, treated with 8% chloroform (v/v), centrifuged, and titered to calculate the plaque-forming units per milliliter (PFU/mL) by serial dilution on an Xcp1 overlay. Phage stocks were cataloged and placed at 4°C in PB for long-term storage.

**Table 1.**
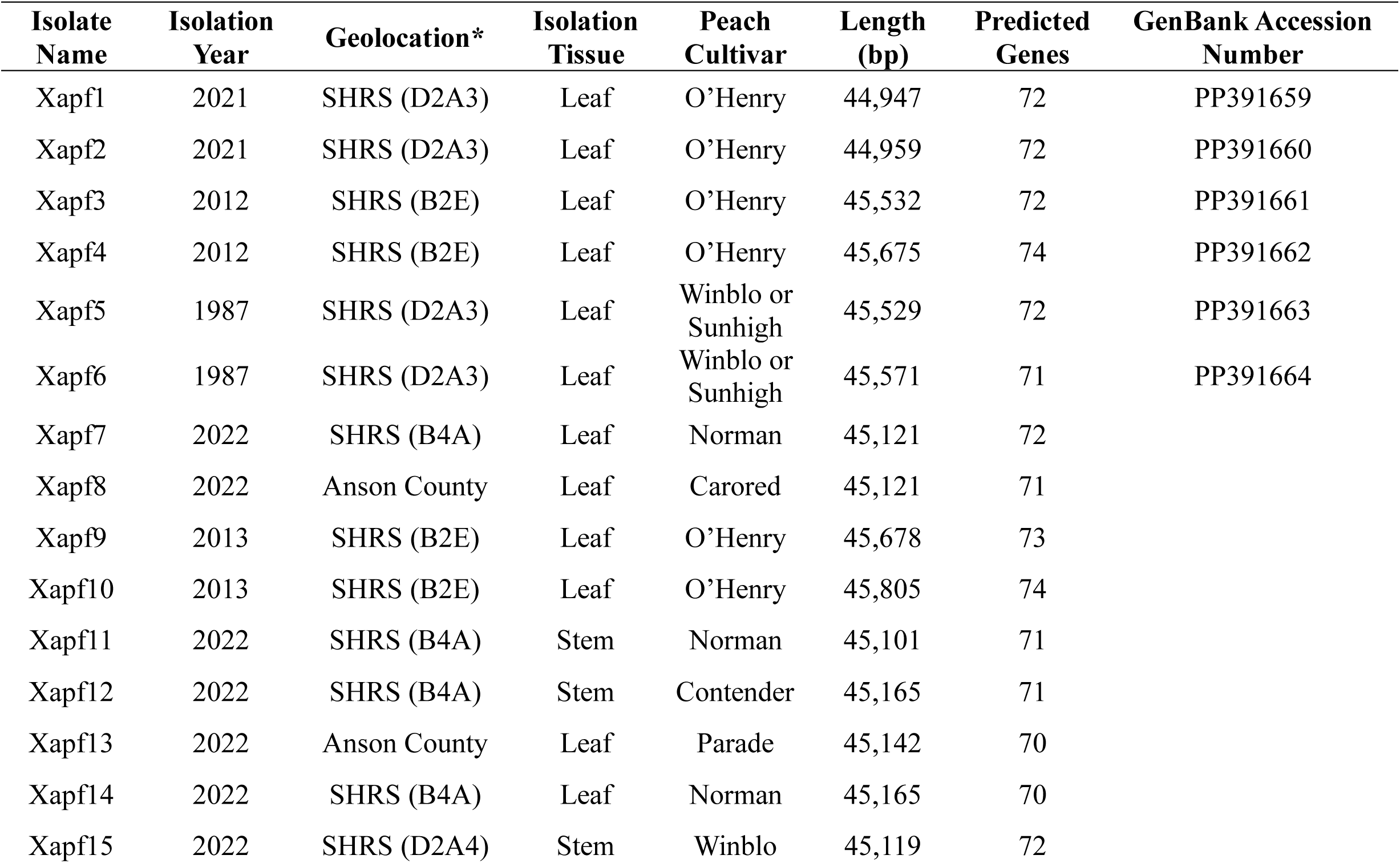
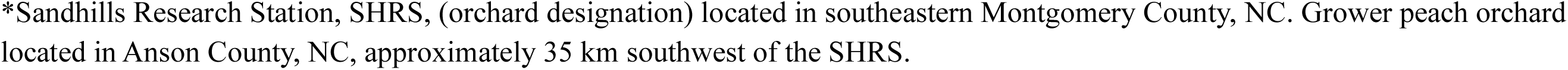
Metadata and genome details for the 15 *Duraznoxanthovirus arenicola* phage isolates described in this geolocation, and tissue of isolation are provided along with the peach cultivar from where the isolates were obtai

| Isolate Name | Isolation Year | Geolocation* | Isolation Tissue | Peach Cultivar | Length (bp) | Predicted Genes | GenBank Accession Number |
| --- | --- | --- | --- | --- | --- | --- | --- |
| Xapf1 | 2021 | SHRS (D2A3) | Leaf | O'Henry | 44,947 | 72 | PP391659 |
| Xapf2 | 2021 | SHRS (D2A3) | Leaf | O'Henry | 44,959 | 72 | PP391660 |
| Xapf3 | 2012 | SHRS (B2E) | Leaf | O'Henry | 45,532 | 72 | PP391661 |
| Xapf4 | 2012 | SHRS (B2E) | Leaf | O'Henry | 45,675 | 74 | PP391662 |
| Xapf5 | 1987 | SHRS (D2A3) | Leaf | Winblo or Sunhigh | 45,529 | 72 | PP391663 |
| Xapf6 | 1987 | SHRS (D2A3) | Leaf | Winblo or Sunhigh | 45,571 | 71 | PP391664 |
| Xapf7 | 2022 | SHRS (B4A) | Leaf | Norman | 45,121 | 72 |  |
| Xapf8 | 2022 | Anson County | Leaf | Carored | 45,121 | 71 |  |
| Xapf9 | 2013 | SHRS (B2E) | Leaf | O'Henry | 45,678 | 73 |  |
| Xapf10 | 2013 | SHRS (B2E) | Leaf | O'Henry | 45,805 | 74 |  |
| Xapf11 | 2022 | SHRS (B4A) | Stem | Norman | 45,101 | 71 |  |
| Xapf12 | 2022 | SHRS (B4A) | Stem | Contender | 45,165 | 71 |  |
| Xapf13 | 2022 | Anson County | Leaf | Parade | 45,142 | 70 |  |
| Xapf14 | 2022 | SHRS (B4A) | Leaf | Norman | 45,165 | 70 |  |
| Xapf15 | 2022 | SHRS (D2A4) | Stem | Winblo | 45,119 | 72 |  |

\*Sandhills Research Station, SHRS, (orchard designation) located in southeastern Montgomery County, NC. Grower peach orchard located in Anson County, NC, approximately 35 km southwest of the SHRS.

### Phage plaque morphology

Double-layer agar plates were prepared with a 30-mL SPA base and a 3-mL top layer of 0.5% water agar seeded with 100 µL of Xap inoculum at 10^6^ CFU/mL. After the inoculated water agar solidified, a 50-µL aliquot of phage suspension (10⁴ PFU/mL) was spread across the top agar layer. After 24 h of incubation at 28 °C, plates were scanned at 600 DPI using an Epson Perfection V600 Photo Scanner. This was repeated twice for each phage isolate.

### DNA extraction

A 2.5-mL aliquot of phage stock in PB (10^9^ PFU/mL) was centrifuged at maximum speed (14,000 rpm) for 3 min. The supernatant was filtered through a 0.22 µm membrane, and the filtrate was transferred to an Amicon® Ultra Centrifugal Filter (100 kDa) and centrifuged at 5,000 rpm for 10 min. The retained concentrate was washed twice with 1 mL of PB, each time centrifuging at 6,000 rpm for 10 min. The purified phage concentrate was transferred to a microfuge tube for DNA extraction using the Phage DNA Isolation Kit (Norgen Biotek Corporation, Thorold, ON, Canada), following the manufacturer’s instructions and including the optional DNaseI and ProteinaseK steps. DNA concentration was measured using the Qubit 4 Fluorometer with the 1× dsDNA High Sensitivity Assay Kit (Thermo Fisher Scientific, USA). Samples were shipped on ice to SeqCenter (Pittsburgh, PA, USA) for sequencing.

### Genome sequencing, assembly, and annotation

Sequencing libraries were prepared by SeqCenter and sequenced on an Illumina NextSeq2000 instrument to produce 2x151 bp reads. Resulting Illumina reads were quality-checked using FastQC v. 0.11.9 and used as input for contig assembly with SPAdes v. 3.15.5 using the *--metaviral* option [19, 20]. The resulting fasta file was analyzed using Virsorter2 v. 2.2.3 with the –*min-length 1500* option to detect viral sequences [21]. Confirmed, viral genomes were stored in individual fasta files and checked for completeness and contamination using CheckV v. 0.9.0 and the CheckV database v. 1.5 [22]. Phage genomes were then annotated using Pharokka v. 1.3.2 [23]. The genome sequences for Xapϕ7-Xapϕ15 were deposited in GenBank under accession numbers: TBD.

### Comparative genomic analyses of Xapϕ phages

The VIRDIC standalone software v. 1.1, with default parameters, was used to calculate intergenic distances of the 15 Xapϕ phages, the output was visualized using the R v. 4.5.0 package pheatmap v. 1.0.12 [24–26]. To examine structural conservation among the 15 Xapϕ phage genomes, all were rotated to start with the *DNA primase* gene. Genomes were then re-annotated with Pharokka v. 1.3.2 to ensure consistent gene predictions and functional assignments. Annotated GenBank files were imported into Clinker v. 0.0.26, and synteny plots were generated using clustermap.js. [27]. In this visualization, parameters were set so that proteins sharing >50% identity were connected by shaded lines of the same color, highlighting conserved regions across genomes. The gene body was colored in black if it belonged to the accessory genome generated as described below. All genes were labeled with their predicted functions if known.

A pangenome for the 15 Xapϕ phages was constructed using custom python scripts for Pharokka, MMSeqs2, MAFFT, trimAI, and IQ-TREE [28–32]. First, the phage protein-coding regions were extracted from the Pharokka annotations. Protein sequences from all phages were merged into a single file and clustered using MMSeqs2 v. 15.6f452 with 70% identity and 50% coverage thresholds [28–30]. A protein presence-absence matrix was created from these clusters using a custom script, *make_pa_matrix.py,* and was used to construct the upset plot using the ComplexUpset v. 1.3.3 package in R [33]. Protein clusters conserved across all 15 phages were aligned with MAFFT in E-INS-i mode and trimmed using trimAI with a gap threshold of 50% (-gt 0.5). A partitioned maximum-likelihood protein phylogeny was calculated using IQ-TREE with ModelFinder to determine the best-fit model and 1000 Ultrafast bootstap and SH-aLRT test replicates [32, 34]. The resulting phylogenetic tree was mid-point rooted using iTOL and visualized using the T-BAS v. 2.4 toolkit on the DeCIFR portal [35, 36].

### Phage genomes selected for comparative analysis

To assess genetic relatedness and diversity, we generated a tblastx distance tree using all ICTV-classified *Caudoviricetes* genomes from the Virus Metadata Resource (MSL40; https://ictv.global/sites/default/files/VMR/VMR_MSL40.v1.20250307.xlsx), along with the nine newly sequenced phages (Xapϕ7–Xapϕ15) from this study. The tree was visualized and annotated in iTOL to identify the *Caudoviricetes* clade containing these Xapϕ phage genomes (Fig. S1) [37]. Genomes phylogenetically closest to Xapϕ7–Xapϕ15 were then selected and downloaded from the NCBI database (Table 1 & S1) [9, 14]. This final dataset comprised 45 genomes: the nine newly sequenced Xapϕ7–Xapϕ15, six previously published Xapϕ phages (Xapϕ1–Xapϕ6), and 30 publicly available genomes.

### Taxonomic classification and comparative genomic analyses of 45 phage genomes

Intergenomic distances among the 45 phages were calculated using the similarity function in taxMyPhage v. 0.3.3 [38]. Protein similarity was assessed with MMSeqs2, clustering proteins at 50% minimum sequence identity and coverage [28]. Protein clusters were extracted as multi-FASTA files, converted into count and presence/absence matrices by genome, and used to calculate Jaccard similarity based on the percentage of shared clusters among the 45 phages. Similarity matrices were visualized using the pheatmap v. 1.0.12 package in R v. 4.5.0 [24].

Phylogenetic relationships were inferred by constructing a pangenome from all 45 genomes using the same methodology applied to the 15 Xapϕ phages. Single-gene phylogenies for the portal protein and terminase large subunit were built using the same parameters as the core phylogeny. Trees were visualized with the T-BAS v. 2.4 toolkit on the DeCIFR portal [35, 36]. Co-phylogenies combining the core partitioned phylogeny and single-gene phylogenies were generated using the ape and phytools packages in R [39].

## Results

### Decoding Xapϕ Phages: Conserved Cores and Divergent Paths Across Decades

*Phenotypic characterization:* A total of nine new phages were isolated, purified, and characterized in this work, named Xapϕ7-Xapϕ15. In addition, six previously characterized Xapϕ phages (Xapϕ1–Xapϕ6) were included in the morphological characterization and genomic analyses used to classify phages virulent on Xap [9]. All 15 Xapϕ phages were lytic against Xap strain Xcp1 in double overlay plate assays (Fig. 1A). Plaque sizes ranged from 250 to 500 µm, with Xapϕ4 producing the smallest and Xapϕ8 the largest plaques. Phages Xapϕ1 and Xapϕ3 exhibited a hazy plaque morphology, whereas Xapϕ7, Xapϕ8, and Xapϕ13 showed large, clear plaques with defined edges. The Xapϕ4, Xapϕ5, Xapϕ6, Xapϕ10, and Xapϕ11 phages showed small pinpoint plaques with a light halo.

**Figure 1.**
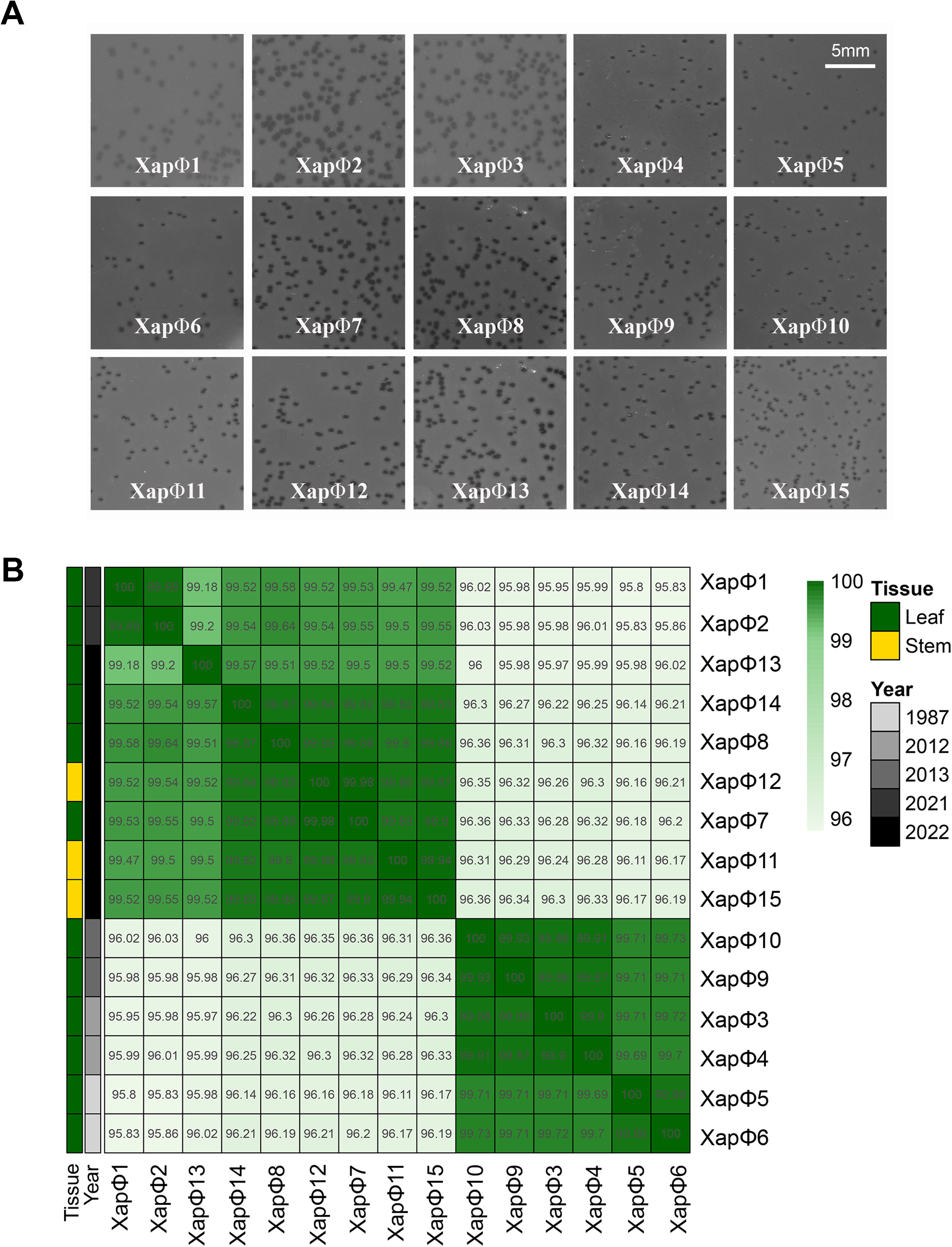
Plaque morphology and genomic relatedness of Xapϕ1–Xapϕ15 phages. (A) Plaque morphology of Xapϕ1–Xapϕ15 phages at a concentration of 10⁴ PFU/mL inoculated onto *Xanthomonas arboricola* pv. *pruni* strain Xcp1 using a double-overlay plaque assay. Images were captured 48 hours post-inoculation with an Epson Perfection V600 Photo Scanner at 600 DPI resolution. A scale bar is shown in the upper-right panel. (B) Heatmap of VIRIDIC-calculated intergenomic distances among Xapϕ1–Xapϕ15 phages, all virulent on *X. arboricola* pv. *pruni* and classified within the newly proposed genus and species, *Duraznoxanthovirus arenicola*. Each cell represents a pairwise intergenomic distances. Tissue, cultivar, and year of isolation are indicated by color coding on the left, with the key provided on the right.

*Genotypic characterization:* Complete genomes were generated for Xapϕ7–Xapϕ15 and their genome sizes ranged from 45,101 bp (Xapϕ11) to 45,805 bp (Xapϕ10), with 70–74 predicted protein-coding sequences per genome (Table 1). To understand the genomic diversity among the 15 Xapϕ phages, which included Xapϕ 7–Xapϕ15 and the six published Xapϕ1–Xapϕ6 genomes, we compared nucleotide similarity by calculating intergenomic distances (Fig. 1B). This revealed that among all 15 genomes, there was >95.8% similarity. The phages clustered into two distinct groups. One group contained Xapϕ3, Xapϕ4, Xapϕ5, Xapϕ6, Xapϕ9, and Xapϕ10, which shared >99.5% similarity, and the second group contained Xapϕ1, Xapϕ2, Xapϕ7, Xapϕ8, Xapϕ11-Xapϕ15, which shared >99.0% similarity (Fig. 1B). Curiously, the phages in the first group were isolated between the years 1987 and 2013. In contrast, the phages in the second group were isolated in 2021 and 2022.

To assess conservation of gene content among the 15 Xapϕ phages, gene synteny was compared using Clinker. This analysis identified conserved genomic regions exhibiting >70% amino acid identity, which included structural components like the tail and capsid proteins, as well as lysis enzymes (Fig. 2A). Despite the high protein conservation among the 15 Xapϕ phages, at least 16 loci showed <70% similarity, including several homing endonucleases, the terminase large subunit, head protein, and several hypothetical proteins (Fig. 2A). These results were supported by a pangenome analysis that elucidated 63 core genes for all 15 Xapϕ phages with >70% identity (Fig. 2B; SFile 1). The terminase large subunit (Xapϕ9: PAQQMYVM_CDS_0028), two homing endonucleases (Xapϕ5: HCUDOGHY_CDS_0062 and Xapϕ9: PAQQMYVM_CDS_0029) and a hypothetical protein (Xapϕ5: HCUDOGHY_CDS_0002) were conserved at >70% similarity among Xapϕ phages isolated between 1987 and 2013 (Xapϕ3- Xapϕ6, Xapϕ9, and Xapϕ10) (SFile 1). Similarly, a second terminase large subunit (Xapϕ12: WIUMOQPX_CDS_0041), a second homing endonuclease (Xapϕ8: MQEUNEBV_CDS_0003), and a hypothetical protein (Xapϕ15: AULOTCPV_CDS_0061) were conserved among Xapϕ phages isolated after 2020 (Xapϕ1, Xapϕ2, Xapϕ7, Xapϕ8, and Xapϕ11-Xapϕ15) (Fig. 2B; SFile 1). These patterns indicate that temporal factors influence phage genome evolution in perennial plant-associated ecosystems.

**Figure 2.**
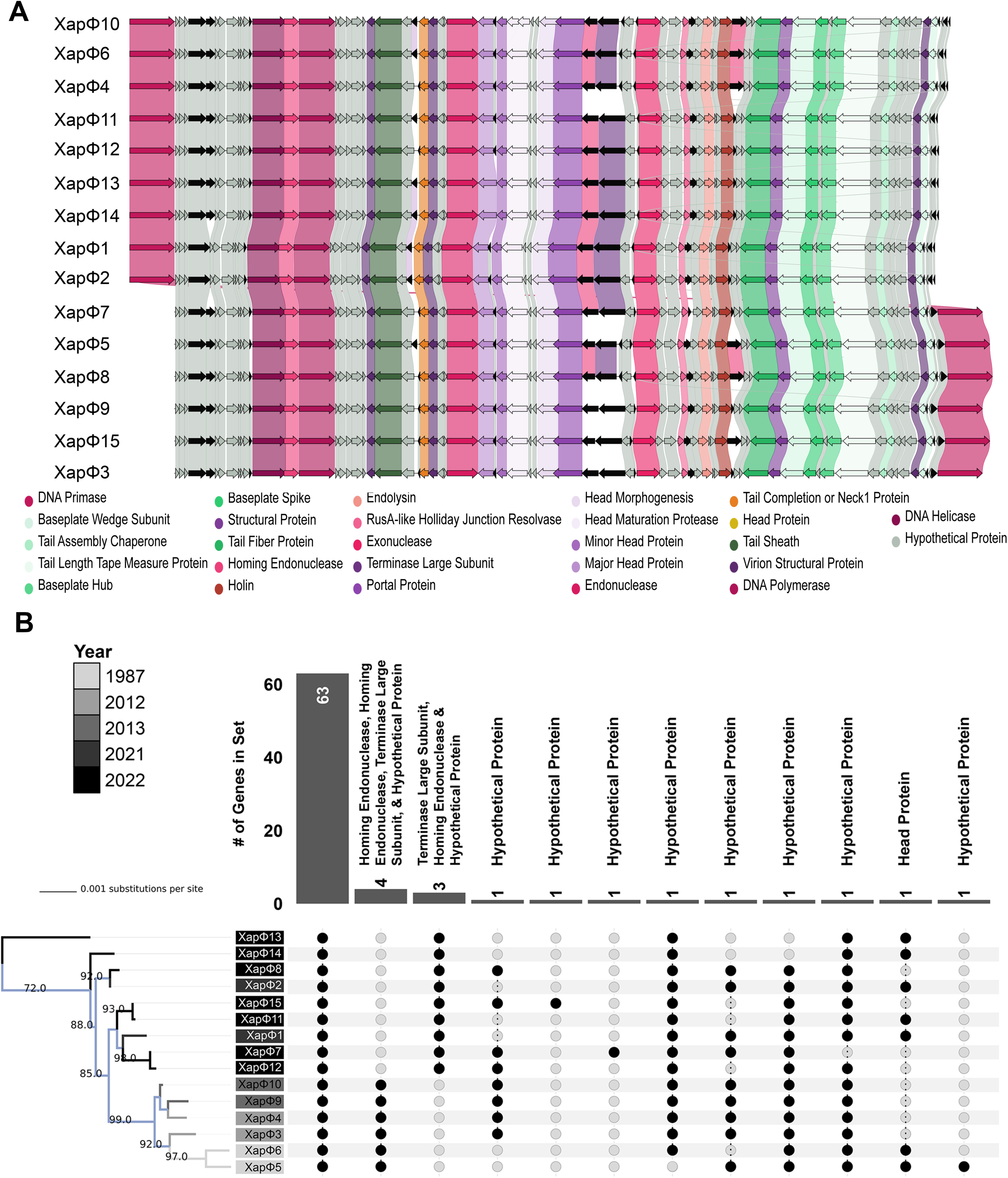
Genomic architecture and evolutionary patterns in *Duraznoxanthovirus arenicola* phages. (A) Genome synteny plot of phages Xapϕ1–Xapϕ15 generated using Clinker. Genes in the core genome are color-coded by predicted function, with hypothetical proteins shown in grey. Genes in the accessory genome are colored in black. Shaded connecting lines indicate >70% amino acid similarity among homologous genes. Functional groups are distinguished by color: green for tail-associated genes, purple for head and packaging, pink for DNA-related functions, and brown for lysis. (B) Midpoint-rooted maximum likelihood phylogenetic tree of Xapϕ1– Xapϕ15, inferred from a core genome alignment using IQ-TREE. Branches and node labels are color-coded by year of isolation. An accompanying Upset plot summarizes pangenome distribution across the phages. The core genome comprises 63 genes, while predicted functions of accessory genes are listed above each gene set.

### Breaking the Mold: Xapϕ Phages Elucidate a New Genomic Frontier for Phages Infecting Plant-Associated Bacteria

*Comparative genomic analysis of Xapϕ phages:* To identify the most closely related phages to the 15 Xapϕ phages, their genomes were imputed into ViPTree to construct a tblastx distance tree (SFig. 1) [40]. Notably, one unclassified phage, NEB7, deposited in GenBank by Omnilytics Inc., clustered within the 15 Xapϕ phages (Fig. 3A). Omnilytics manufactures phage-based therapeutics for managing bacterial pathogens in agricultural settings and has a specific product for Xap named AgriPhage-Nut and Stone Fruit Biologicals (https://agriphage.com/product-info/agriphage-nut-stone-fruit-peach-bacterial-spot/). The 15 Xapϕ phages with NEB7 formed a monophyletic clade within the class *Caudoviricetes* and the existing virus subfamily *Kantovirinae*, confirming that these isolates are tailed double-stranded DNA lytic phages [9]. Within *Kantovirinae*, there are two validly named genera: *Beograduvirus* (four isolates) and (13 isolates), both of which house phages that infect bacterial plant pathogens in the genus *Xanthomonas* (Fig. 3A) (MSL40; https://ictv.global/sites/default/files/VMR/VMR_MSL40.v1.20250307.xlsx) [41]. However, the Xapϕ phages constitute a distinct monophyletic clade within the *Kantovirinae* that is closely related to, but separate from, all currently classified genera and species, which we name *Duraznoxanthovirus arenicola*.

**Figure 3.**
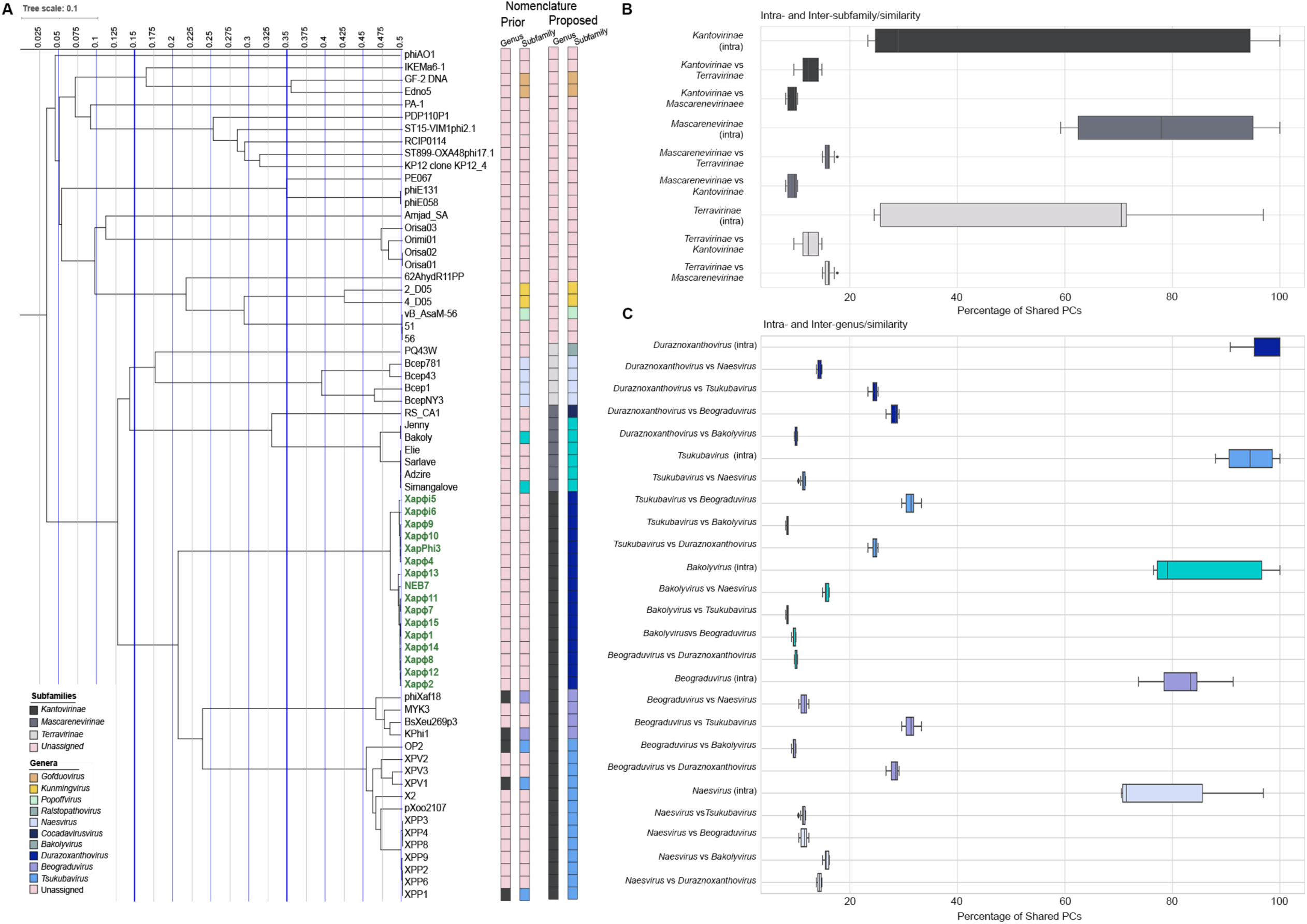
Comparative proteomic relationships among phage subfamilies and genera in *Phytobacteriaviridae*. (A) Phylogenetic tree illustrating evolutionary relationships among phage genera within the subfamilies ***Kantovirinae, Mascarenevirinae***, and ***Terravirinae*** of the newly described family *Phytobacteriaviridae*. Bars to the right indicate prior and proposed nomenclature. The tree scale represents 0.1 substitutions per site. (B) Boxplots showing intra- and inter-subfamily protein similarities among ***Kantovirinae****, **Mascarenevirinae***, and ***Terravirinae***. (C) Boxplots showing intra- and inter-genus protein similarities including *Tsukubavirus*, *Bakolyvirus*, *Duraznoxanthovirus*, *Beograduvirus*, and *Naesvirus*. Protein similiarity is based on the percentage of shared protein clusters (PCs) among and between taxanomic units. The x-axis indicates the percentage of shared PCs, and the y-axis represents intra- or inter-paired comparisons at the subfamily (B) or genus (C) level.

The discovery and naming of *Duraznoxanthovirus arenicola* led to the identification of a new monophyletic clade adjacent to *Kantovirinae* and *Mascarenevirinae* in the VipTree analysis, which we designate as *Terravirinae* (Fig. 3A, 3B). This clade was confirmed as a distinct subfamily through shared protein cluster (PC) analysis, VIRIDIC intergenomic distance calculations, and Jaccard similarity indices (Fig. 3B; SFig. 2A, 2B). Intra-subfamily comparisons showed >20% shared PCs. In contrast, inter-subfamily comparisons among members of the three subfamilies within the family *Anamaviridae* were substantially lower (Fig. 3B). Additionally, VIRIDIC and Jaccard analyses revealed >32.3% identity and >23.4% similarity among genera in the *Kantovirinae*; >67.1% identity and >59.2% similarity among genera in the *Mascarenevirinae*; and >33.5% identity and >24.5% similarity among genera in the *Terravirinae* (SFig. 2A, 2B). Together, these results introduce a previously unrecognized lineage and formalize *Terravirinae* as a new subfamily within the recently elucidated phage family *Anamaviridae* [14].

Furthermore, ViPTree analysis indicates that within *Terravirinae* there are two genera: the validly named *Naesvirus* and a newly identified clade represented by *Ralstopathovirus humicola* (PQ43W) (Fig. 3A; SFig. 1B) [42]. The delineation of this new genus and species within *Terravirinae*, as distinct from the recently described subfamily *Mascarenevirinae* in the family *Anamaviridae*, is strongly supported by multiple metrics, including high intra-versus inter-genus shared PC similarity, VIRIDIC intergenomic distances, and Jaccard similarity indices (Fig. 3C; SFig. 2A, 2B). Specifically, intra-genus comparisons showed >70% shared PC, whereas inter-genus comparisons among phage genera in the *Anamaviridae* were substantially lower (Fig. 3C). Furthermore, *Naesvirus* and *Ralstopathovirus* shared <35% intergenomic distance and <25.7% Jaccard similarity with other genera in this family (Fig. 3C; SFig. 2A, 2B).

To identify shared and unique genes across phage families, subfamilies, and genera: *Anamaviridae*; *Mascarenevirinae*, *Terravirinae*, and *Kantovirinae*; *Cocadavirus*, *Bakolyvirus*, *Naesvirus*, *Ralstopathovirus*, *Duraznoxanthovirus*, *Beograduvirus*, and *Tsukubavirus*, we performed a pangenome analysis with the 45 phages that comprise this clade (Fig. 4A; SFile2). The *Anamaviridae* core genome consisted of ten genes, where each gene shared >70% protein similarity, including the tail completion or Neck1 protein, portal protein, terminase large subunit, baseplate wedge subunit, DNA helicase, tail sheath protein, virion structural protein, DNA polymerase, and two hypothetical proteins (Fig. 4A; SFile2). At the subfamily level, there were 26 core genes unique to phages in the *Mascarenevirinae*, six core genes in the *Terravirinae*, and 12 unique to the *Kantovirinae* subfamily (Fig. 4A; SFile 2). At the genus level, there were 13 core genes unique to the *Cocadavirus,* 35 to the *Bakolyvirus,* 25 to the *Naesvirus,* 38 to the *Ralstopathovirus*, 28 to the *Duraznoxanthovirus*, 20 to the *Beograduvirus*, and 31 to the *Tsukubavirus* (Fig. 4A; SFile 2). Interestingly, the genus with the fewest core genes (*Cocadavirus*) has only one member, phage RS_CA1 [14]. Similarly, the genus with the most core genes (*Ralstopathovirus*) also has only one member, phage PQ43W [42]. This hierarchical distribution of core genes shared among phage genomes within the *Anamaviridae* and across its subfamilies and genera underscores precise evolutionary trajectories that align with the bacterial hosts they infect (Fig. 4A).

**Figure 4.**
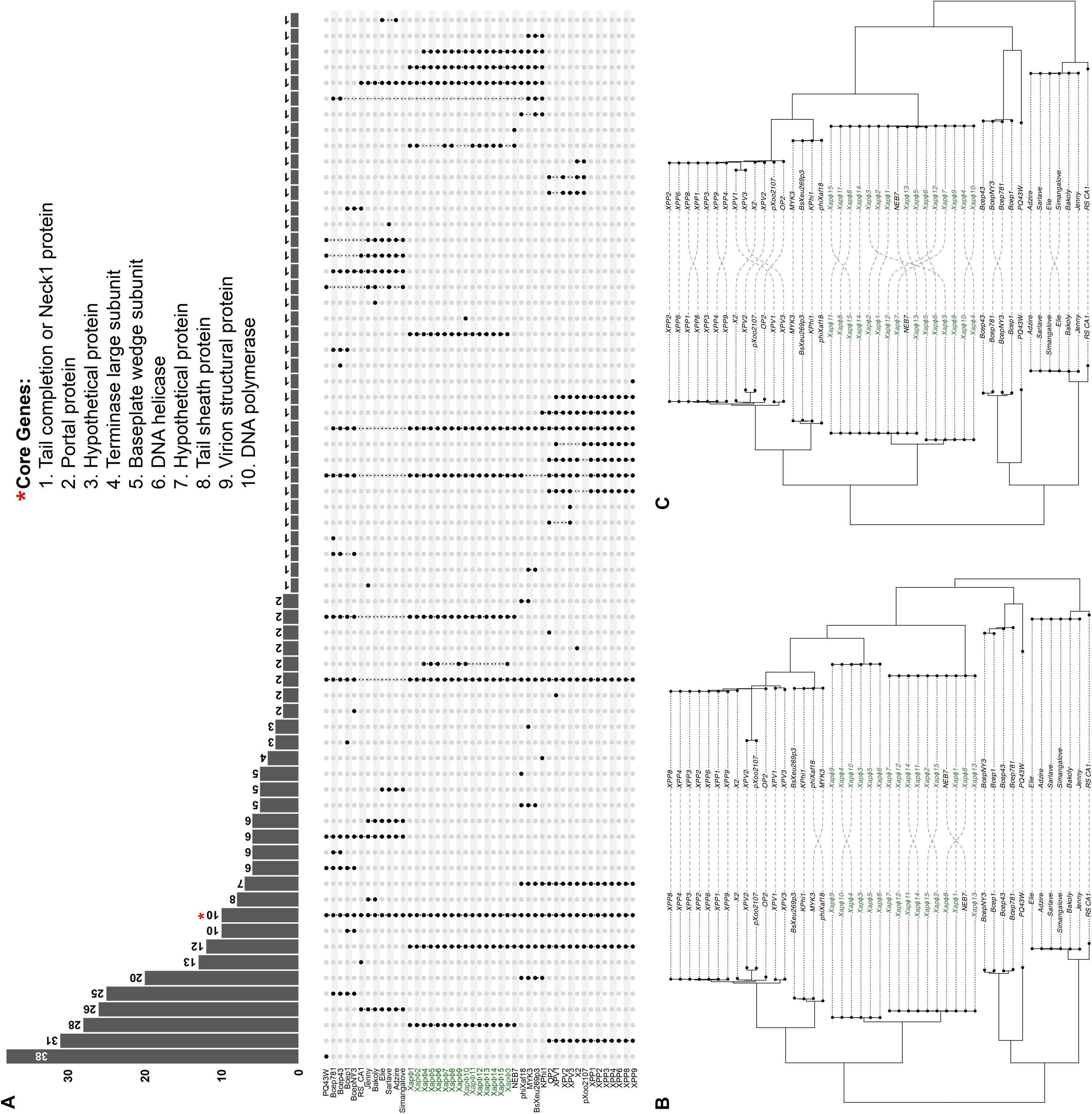
Genomic connectivity: Pangenome composition and phylogenetic concordance of *Phytobacteriaviridae.* (A) Upset plot summarizing the pangenome distribution of Xapϕ1– Xapϕ15 and 30 closely related ***Phytobacteriaviridae*** phages. The set of 10 core genes is marked with a red star, and their predicted function is embedded in the plot. (B & C) Tanglegram illustrating the correspondence between the core genome maximum likelihood phylogenetic tree of all ***Phytobacteriaviridae*** phages (left) and the conserved terminase large subunit proteins (right) (B) or conserved phage portal proteins (right) (C).

Beyond the conserved core, lineage-specific genes highlight functional and ecological differences among genera within *Anamaviridae*. For example, seven genes were unique to phages in the *Beograduvirus* and *Tsukubavirus*, which infect Xanthomonads pathogenic on tomato and pepper, and rice, respectively, but are absent in *Duraznoxanthovirus arenicola* phages that infect Xanthomonads of peach. Two genes encoding DNA primase (subfamily_134) and baseplate wedge subunit (subfamily_48) were conserved across all phages in *Anamaviridae* except for the seven phages in the subfamily *Mascarenevirinae*, all of which infect *Ralstonia* (Fig. 4A; SFig. 2; SFile 2). Two additional genes which encode tail length tape measure protein (subfamily_183) and a hypothetical protein (subfamily_181) were conserved in all *Duraznoxanthovirus, Naesvirus*, and *Ralstophatovirus* phages, which infect Xanthomonads on peach, *Burkholderia*, and *Ralstonia*, respectively, but were absent in the *Tsukubavirus* (which infects *Xanthomonas* pathogenic on rice), *Beograduvirus* (infecting *Xanthomonas* pathogenic on tomato and pepper), *Bakolyvirus* (which infect *Ralstonia*), and *Cocadavirus* (which infect *Ralstonia*) (Fig 4A; SFile 2). Only one gene encoding a hypothetical protein (subfamily_187) was unique to the 15 Xapϕ phages, excluding NEB7 (SFile 2).

Finally, a tanglegram phylogenetic tree inferred from the concatenated alignment of ten core genes and the TerL or Portal protein resolved highly supported monophyletic clades for the subfamilies and genera comprising this family of phages virulent on plant-associated bacterial hosts (Fig. 4B, 4C). There was strong phylogenetic congruence between the core-gene phylogeny (>98% bootstrap support) and those generated from the two marker genes. Specifically, the 15 Xapϕ phages and NEB7 formed a congruent monophyletic clade encompassing *Duraznoxanthovirus arenicola.* This clade along with four confirmed phages from the genus *Beogradivirus* and 13 confirmed phages from genus *Tsukubavirus*, comprised the subfamily *Kantovirniae*. Adjacent to this group was the *Terravirinae* subfamily, composed of four *Naesvirus* phages and one *Ralstopathovirus* phage. Lastly, the subfamily *Mascarenevirnae* was supported by a clade of four *Bakolyvirus* phages and one *Cocadavirus* phage (Fig. 4B, 4C). The intact clades across both trees indicate robust evolutionary relationships within *Anamaviridae*, reinforcing the proposed taxonomic framework and highlighting deep phylogenetic structure among the phage subfamilies and genera.

## Discussion

### Genomic stability meets subtle divergence: Insights from four decades of Xapϕ phages

A central challenge in advancing phage biology in plant-associated systems is the accurate classification and naming of organisms, as diversity in these ecosystems remains poorly understood. In this study, we conducted a comprehensive comparative genomic and proteomic analysis of 15 Xapϕ phages virulent on Xap, the causal agent of bacterial spot of *Prunus* species. Interestingly, these 15 isolates segregate into two well-supported clades that correspond to their temporal origin: one consisting of phages recovered between 1987 and 2013 (>99.7% identity), and the other comprising isolates obtained after 2020 (>99.2% identity). This divergence among the 15 Xapϕ phages suggests a minimal yet impactful biotic/abiotic perturbation in the Sandhills region of North Carolina between 2013 and 2020, which may have contributed to observed phage evolution.

Here, we reveal that *Duraznoxanthovirus arenicola* exhibits remarkable genomic conservation, maintaining over 95% nucleotide identity across isolates collected over nearly four decades from the Sandhills regions in North Carolina, the historically (since the early 20^th^ century) central peach production region of the state. This long-term stability underscores strong purifying selection acting on 63 essential genes encoding proteins critical for DNA packaging and virion architecture. However, this conservation is not absolute. Localized variability in accessory loci, including the terminase large subunit, two homing endonucleases, and numerous hypothetical proteins, suggests adaptive fine-tuning to the bacterial host within peach production ecosystems. These genes, often associated with recombination and host interactions, may drive microevolutionary shifts that enable persistence under fluctuating biotic and abiotic conditions. Importantly, all phages were isolated using a single bait strain of Xap (Xcp1, isolated in 1984), which may have constrained observed diversity and influenced inferred evolutionary trajectories [15]. Nonetheless, the coexistence of deep genomic stability among phages with targeted diversification of specific genes reflects a dual evolutionary strategy that preserves core functionality while exploring adaptive niches in a dynamic perennial agricultural environment.

Contrary to expectations of extensive genomic mosaicism across temporal scales, these patterns suggest that vertical evolutionary processes may dominate in plant-associated perennial ecosystems. This challenges the prevailing idea that phage genomes are inherently dynamic and fragmented over time [49, 50]. Instead, the strong genomic conservation observed among Xapϕ phages across nearly four decades, all isolated from a single geographic region, indicates that stable ecological conditions and persistent host–phage relationships limit diversification despite temporal separation [51]. This reframes our understanding of phage evolution in perennial agricultural systems, emphasizing how ecological stability can favor lineage continuity over genomic flux. These findings are highly encouraging for therapeutic applications, as they suggest predictable performance and a lower risk of genome instability in phage-based management strategies. However, a critical question remains: *If genomic stability is so pronounced, why do phage-based treatments in agricultural systems often produce inconsistent results?*

### Genomic clues to host specificity: Insights from *Kantovirinae* phages

Phages with narrow host ranges often specialize and contribute to long-term ecological stability within specific niches [43–46]. These phages, commonly referred to as *monovalent*, typically exhibit lytic activity restricted to bacterial species or strains within a single genus [47]. Based on our genomic analysis, we hypothesize that all phages within the *Kantovirinae* subfamily are monovalent to *Xanthomonas*. The 12 core genes shared among phages in this subfamily could be responsible for delineating this host range. Furthermore, a finer host range resolution exists among genera within the *Kantovirinae* subfamily, where *Duraznoxanthovirus* are lytic on *X. arboricola* pv. *pruni*; *Tsukubavirus* are lytic on *X. oryzae* pv. *oryzae*; and *Beograduvirus* are lytic on *Xanthomonas* infecting pepper and tomato.

In contrast, polyvalent phages display broader host ranges and can lyse bacteria across multiple genera [48]. Although lytic phenotypes were not evaluated in this study, it is plausible that some members within and among the *Anamaviridae* and *Kantovirinae* may infect hosts from more than one bacterial genera and/or species. Confirming this hypothesis will require comprehensive phenotypic assays on individual phage and bacterial isolates from different viral and bacterial genera and species to elucidate the key factors driving monovalent and polyvalent interactions in plant-associated ecosystems.

### *Anamaviridae*: A family of phages infecting plant-associated bacteria

Phages are among the most abundant biological entities on Earth, with estimates suggesting roughly ten phage particles for every bacterial cell [1]. Yet, despite this ubiquity, their diversity and ecological roles in plant-associated environments remain poorly characterized, particularly for phages infecting bacterial plant pathogens in agricultural systems. This gap constrains efforts to harness phages as biocontrol agents for sustainable disease management. Addressing this challenge begins with defining the phages that exist, their bacterial hosts, and the evolutionary dynamics that shape these interactions [3].

Beyond characterizing a new genus and species, *Duraznoxanthovirus arenicola*, within the *Kantovirinae* subfamily, our analyses support a broader taxonomic restructuring of the recently delineated lineage of phages within the class *Caudoviricetes*, family *Anamaviridae*, which encompasses phages virulent on bacterial hosts associated with plants [14, 49]. Within this family, three distinct subfamilies are now resolved based on phylogenetic clustering, intergenomic distances, and shared protein content: *Kantovirinae*, *Mascarenevirinae*, and *Terravirinae.* The subfamily *Kantovirinae* comprises the genera *Duraznoxanthovirus*, *Beograduvirus,* and *Tsukubavirus*, all of which infect plant-pathogenic *Xanthomonas* species [50]. The subfamily *Mascarenevirinae* comprises *Bakolyvirus* and *Cocadavirus*, which target plant-pathogenic *Ralstonia* species [51]. The subfamily *Terravirinae* includes *Naesvirus* and *Ralstopathovirus,* each associated with bacterial hosts in the genera *Burkholderia* and *Ralstonia,* respectively [42, 52]. This hierarchical framework reflects the deep evolutionary divergence among phages infecting plant-associated bacteria and provides a foundation for future ecological studies within agricultural microbiomes (Huerta, *submitted*). The congruence between phylogenies inferred from core genes and marker proteins further reinforces the robustness of this classification. Biological validation remains essential to determine the functional relevance of the observed genomic diversity within these recently elucidated phage subfamilies and genera.

The proposed taxonomic framework not only organizes existing diversity but also provides a predictive structure for future discoveries. Each subfamily exhibits distinct evolutionary trajectories aligned with their bacterial hosts: *Kantovirinae* phages specialize on *Xanthomonas* species, while *Mascarenevirinae* and *Terravirinae* primarily target *Ralstonia* and *Burkholderia* genera, respectively [14]. The genera within these subfamilies, *Duraznoxanthovirus*, *Beograduvirus*, *Tsukubavirus*, *Bakolyvirus*, *Ralstopathovirus*, *Naesvirus*, and *Cocadavirus*, represent deeply divergent lineages supported by phylogenetic clustering, intergenomic distances, and shared protein content. These results underscore the ecological and evolutionary significance of host specialization and highlight the need for expanded phage sampling across plant-associated microbiomes to refine genus-level boundaries and uncover additional diversity. Ultimately, this taxonomic framework lays the foundation for integrating phage ecology with applied strategies for sustainable disease management in agriculture. It may also help inform regulatory practices as we expand the use of phage-based biologicals in agricultural systems.

## Supporting information

SFile 1

SFile 2

## Acknowledgements

We would like to acknowledge the staff at the Sandhills Research Station (Jackson Springs, NC, USA), especially station manager Jeremy Martin, for their assistance in orchard maintenance for many years. This work was supported by the Extension Capacity Fund (Smith-Lever 3(b) and 3(c)), project award no. 7007436, from the U.S. Department of Agriculture’s National Institute of Food and Agriculture, the Genetics and Genomics Institute at North Carolina State University, and the North Carolina Agricultural Research Service.

## Statements and Declarations

The authors have no relevant financial or non-financial interests to disclose.

## Author Contributions

All authors contributed to the conception and design of the study. David F. Ritchie established the peach research orchards, collected the pre-2020 samples, and isolated bacteriophages from those samples. Material preparation, data collection, and post 2020 samples were performed and collected by Katherine M. D. Willman, Meaghan Flaherty and Alejandra Huerta. Katherine M. D. Willman, Prasanna Joglekar, and Dann Turner performed data analysis. Katherine M. D. Willman, Prasanna Joglekar, and Alejandra Huerta prepared visualizations. The first draft of the manuscript was written by Katherine M. D. Willman, Alejandra Huerta, and Meaghan Flaherty, and all authors commented on previous versions of the manuscript. All authors read and approved the final manuscript.

## Data Availability

The datasets generated during and/or analyzed during the current study are available in the NCBI GenBank repository under the following accession numbers: TBD. NCBI GenBank submission IDs: 3011585, 3011590, 3011599, 3011615, 3012150, 3012927, 3012930, 3012932, and 3012933. All scripts and their usage are available on <u>GitHub</u>.

