## Supplemental Material for "Four Decades of Genomic Stability and Adaptive Divergence in *Xanthomonas arboricola* pv. *pruni*-Infecting Phages: Defining *Duraznoxanthovirus arenicola* and Its Evolutionary Framework"

**Table 1.** Summary of the 30 publicly available *Phytobacteriaviridae* phage genomes as of October 27th, 2025.

| Phage | Bacterial Host | Phage Genus | Length (bp) | NCBI GenBank Accession Number |
|---|---|---|---|---|
| Bcep1 | *Burkholderia cepacia* | *Naesvirus* | 48,177 | NC_005263.2 |
| Bcep43 | *Burkholderia cepacia* | *Naesvirus* | 48,024 | NC_005342.2 |
| Bcep781 | *Burkholderia cepacia* | *Naesvirus* | 48,247 | NC_004333.2 |
| BcepNY3 | *Burkholderia cenocepacia* | *Naesvirus* | 47,382 | NC_009604.1 |
| Adzire | *Ralstonia pseudosolanacearum* | *Bakolyvirus* | 44,732 | NC_054462.1 |
| Bakoly | *Ralstonia pseudosolanacearum* | *Bakolyvirus* | 44,222 | NC_054463.1 |
| Elie | *Ralstonia pseudosolanacearum* | *Bakolyvirus* | 44,966 | MT740735.1 |
| Jenny | *Ralstonia pseudosolanacearum* | *Bakolyvirus* | 43,921 | MT740744.1 |
| Sarlave | *Ralstonia pseudosolanacearum* | *Bakolyvirus* | 44,858 | MT740746.1 |
| Simangalove | *Ralstonia pseudosolanacearum* | *Bakolyvirus* | 44,834 | NC_054946.1 |
| RS_CA1 | *Ralstonia solanacearum* & *Ralstonia pseudosolanacearum* | Unclassified | 46,254 | PP316169.1 |
| PQ43W | *Ralstonia pseudosolanacearum* | Unclassified | 47,156 | PP405626.1 |
| BsXeu269p3 | *Xanthomonas euvesicatoria* | *Beograduvirus* | 46,280 | ON996340.1 |
| KPhi1 | *Xanthomonas euvesicatoria* | *Beograduvirus* | 46,077 | NC_054460.1 |
| MYK3 | *Xanthomonas* spp. | *Beograduvirus* | 47,500 | OK275494.1 |
| phiXaf18 | *Xanthomonas vesicatoria* | *Beograduvirus* | 47,407 | NC_054461.1 |
| pXoo2107 | *Xanthomonas oryzae* pv. *oryzae* | *Tsukubavirus* | 46,052 | OP067662.1 |
| X2 | *Xanthomonas oryzae* pv. *oryzae* | *Tsukubavirus* | 45,966 | MW435566.1 |
| OP2 | *Xanthomonas oryzae* pv. *oryzae* | *Tsukubavirus* | 46,643 | NC_007710.1 |

| | | | | |
|---|---|---|---|---|
| XPV1 | *Xanthomonas oryzae* pv. *oryzae* | *Tsukubavirus* | 46,503 | NC_054459.1 |
| XPV2 | *Xanthomonas oryzae* pv. *oryzae* | *Tsukubavirus* | 45,969 | MG944235.1 |
| XPV3 | *Xanthomonas oryzae* pv. *oryzae* | *Tsukubavirus* | 47,046 | MG944236.1 |
| XPP1 | *Xanthomonas oryzae* pv. *oryzae* | *Tsukubavirus* | 46,195 | NC_054458.1 |
| XPP2 | *Xanthomonas oryzae* pv. *oryzae* | *Tsukubavirus* | 46,480 | MG944228.1 |
| XPP3 | *Xanthomonas oryzae* pv. *oryzae* | *Tsukubavirus* | 49,612 | MG944229.1 |
| XPP4 | *Xanthomonas oryzae* pv. *oryzae* | *Tsukubavirus* | 47,397 | MG944230.1 |
| XPP6 | *Xanthomonas oryzae* pv. *oryzae* | *Tsukubavirus* | 46,281 | MG944231.1 |
| XPP8 | *Xanthomonas oryzae* pv. *oryzae* | *Tsukubavirus* | 46,278 | MG944232.1 |
| XPP9 | *Xanthomonas oryzae* pv. *oryzae* | *Tsukubavirus* | 48,669 | MG944233.1 |
| NEB7 | *Xanthomonas* spp. | Unclassified | 45,241 | OQ676962.1 |

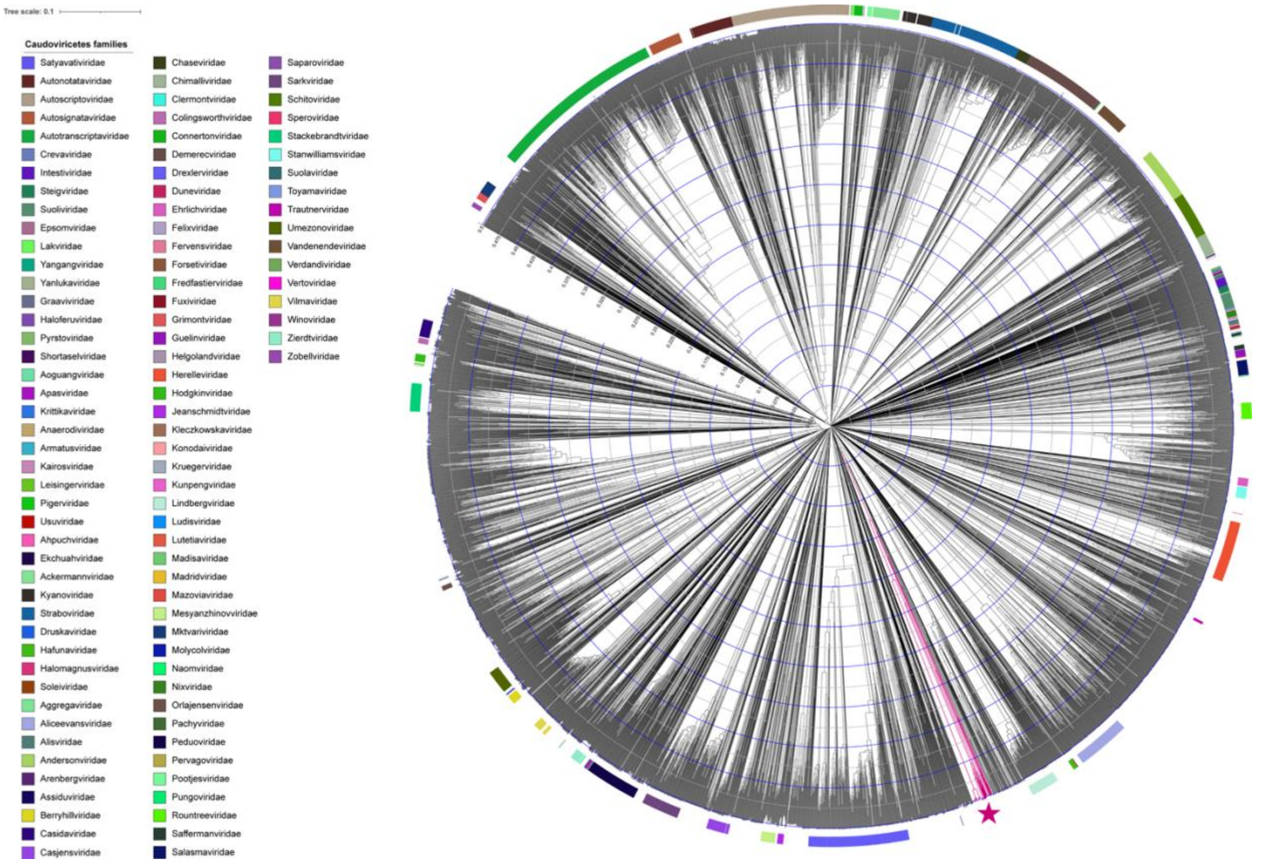


**Figure S1.** Viral proteomic tree (ViPTree) constructed from 5,822 reference genomes, with the 15 Xapϕ phages infecting *Xanthomonas arboricola* pv. *pruni* (Xap) highlighted in magenta and marked with a star. Taxonomic family assignments for reference genomes are indicated by the color of the outer bar.

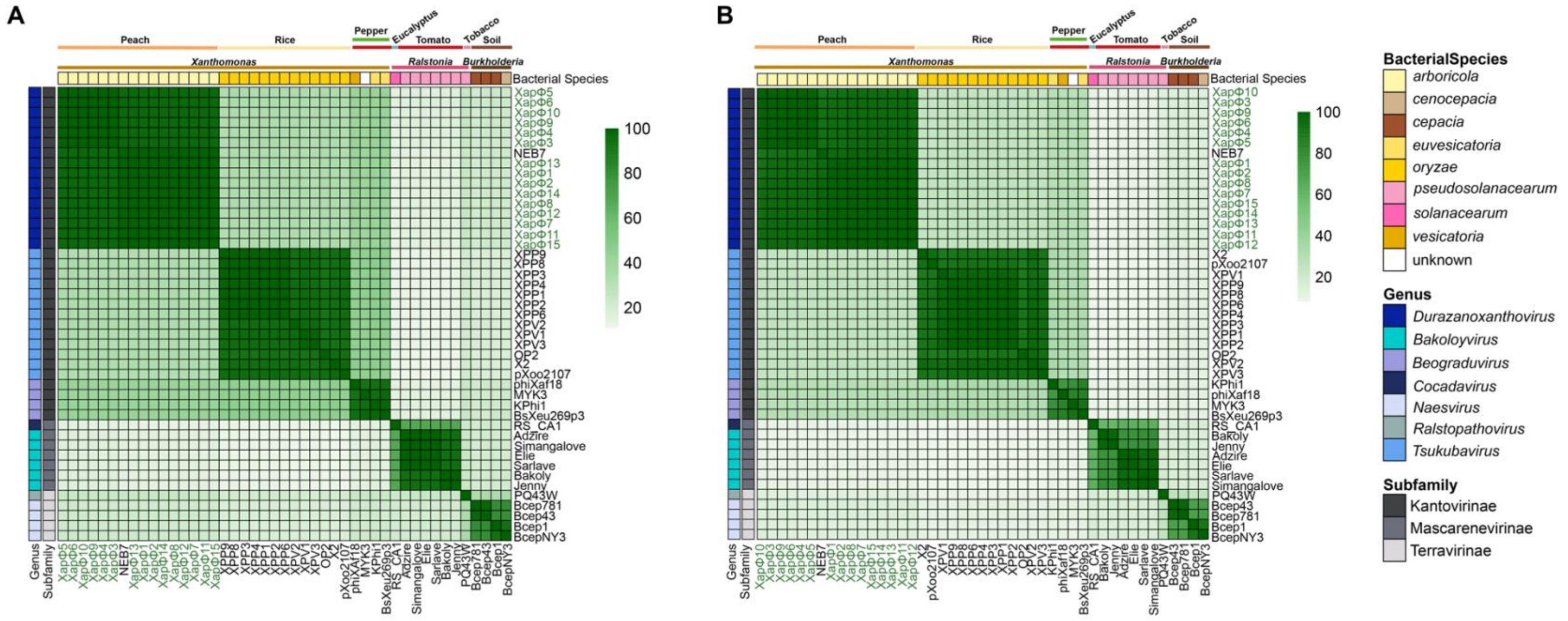


**Figure S2.** Genomic and proteomic similarity analyses reveal a novel phage clade: *Duraznoxanthovirus arenicola*. (A) Heatmap of VIRIDIC-calculated intergenomic distances among Xapϕ1–Xapϕ15 and 30 phage genomes in the *Anamaviridae* family. The Xapϕ1–Xapϕ15 cluster, in conjunction with phage NEB7, forms a distinct, *Duraznoxanthovirus arenicola*. (B) Heatmap of Jaccard similarity index values based on protein identity shared among Xapϕ1–Xapϕ15 and 30 phage genomes in the *Anamaviridae*. Phage genus and subfamily are color-coded as indicated by the bar on the left, with the key on the right. The bacterial host genus, species, and isolation source are listed at the top of each heatmap.

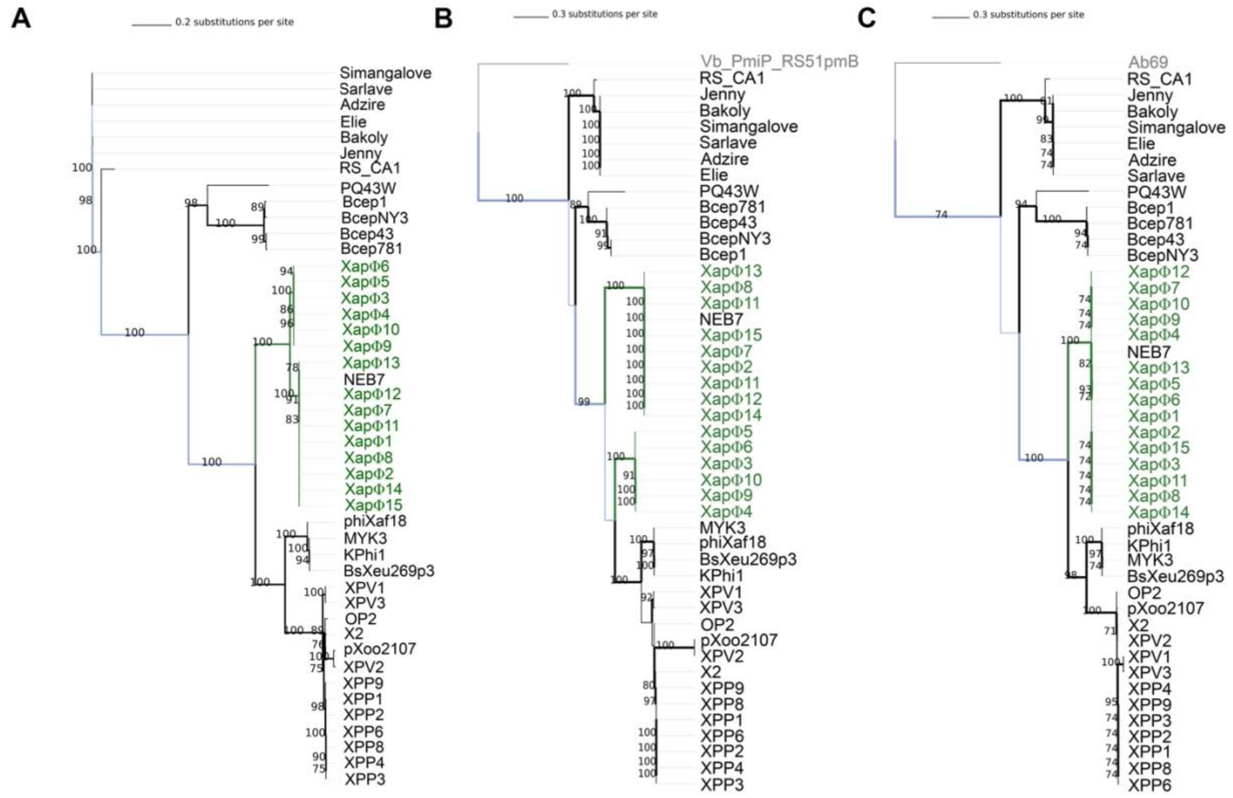


**Figure S3.** Phylogenetic analyses of *Duraznoxanthovirus arenicola* phages reveal evolutionary relationships within the *Anamaviridae*. (A) Midpoint-rooted maximum likelihood phylogenetic tree of Xapϕ1–Xapϕ15 and 30 closely related *Anamaviridae* phages, inferred from a core genome alignment of ten conserved genes using IQ-TREE. Maximum likelihood phylogenetic tree of Xapϕ1–Xapϕ15 and 30 closely related *Anamaviridae* phages, inferred from an amino acid alignment of the (B) TerL sequence and (C) the portal protein sequence using IQ-TREE. The TerL sequence from *Proteus* phage vB_PmiP_RS51pmB (QDH85558.1) and the portal protein sequence from *Acinetobacter* phage Ab69 (WMC00310.1) were used as outgroups, respectively.
